# Maternal diet exerts sex-specific effects on offspring’ personalities in predatory mites

**DOI:** 10.1101/2025.06.13.659584

**Authors:** Thi Hanh Nguyen, Peter Schausberger

**Affiliations:** Department of Behavioral and Cognitive Biology, University of Vienna, Djerassiplatz 1, 1030 Vienna, Austria

## Abstract

Animal personality is characterized by consistent behavior within individuals linked to consistently variable behavior among individuals in a population across time and contexts. Genetic determination, transgenerational effects, and personal experience are major pathways shaping animal personalities. Among these pathways, little attention has been paid to environmental factors in the parental generation affecting offspring personality. Here we tested the effects of the maternal diet on offspring personality in the plant-inhabiting predatory mite Amblyseius swirskii. Mated females and males, whose mothers were fed during egg production on either cattail pollen, two-spotted spider mites, or thrips, were subjected to a battery of three to five tests each for exploration, activity, and boldness. Movement activity was assessed in the mites’ familiar environment. Exploration was quantified by the latency to leave and reach novel sites or objects. Boldness was evaluated by residence in risky and benign sites. Mean behaviors were analyzed by generalized estimating equations, repeatability was assessed by intraclass correlation coefficients. On average, offspring from spider mite-fed mothers were the most active and those from pollen-fed mothers were the shiest. Offspring from thrips-fed mothers were more repeatable in activity than offspring from pollen-and spider mite-fed mothers. Consistently little and highly active personalities produced more eggs than inconsistent, flexible types. Only offspring from pollen-fed mothers were repeatable in boldness. Maternal diet did not influence offspring’ personality in exploration. Taken together, our study suggests that the maternal diet critically influences both mean behavioral trait expression and behavioral repeatability of offspring. The ability of mothers to respond to short-term diet changes during internal egg production allows to adaptively adjust the behavior and personalities of daughters and sons within local and regional groups.

## Introduction

Animal personality refers to behaviors that are consistent within individuals but consistently variable among individuals in a population across time and contexts (Réale et al., 2007). Activity, exploration, aggressiveness, boldness and sociability are the five behavioral dimensions widely used to describe animal personalities (Réale et al., 2007). Environmental conditions, along with genetic architecture can mediate the expression of animal personality (Réale et al., 2007; Dingemanse & Dochtermann, 2013). This applies particularly to environmental conditions experienced during the early phases of life but also to those experienced by parents and passed on to offspring (Stamps & Groothuis, 2010; Reddon, 2012; Sih et al., 2015; Bell & Hellmann, 2019; Stamps & Bell, 2021). Such effects often come from variation in predation pressure (Kortet et al., 2015; Si et al., 2023; Schausberger et al., 2024), food availability and/or nutritional quality (Sebastien et al., 2016; Han & Dingemanse, 2015; Herath et al., 2021; Ersoy et al., 2022; Nguyen & Schausberger, 2024), or social conditions (Jolles et al., 2016; Zajitschek et al., 2017; Horváth et al., 2019; Skinner et al., 2022; Schausberger & Nguyen, 2024). The effects of personal early life experiences on the formation of animal personality have been documented in various taxa, including vertebrates and invertebrates (snails: Tariel et al., 2020; tropical skinks: de Jong et al., 2022; leaf beetles: Tremmel & Müller, 2013; predatory mites: Nguyen & Schausberger, 2024; Schausberger & Nguyen, 2024; Schausberger et al., 2024; Nguyen & Schausberger, 2025). In contrast, parental effects on animal personalities have not been well researched (Reddon, 2012; Zajitschek et al., 2017; Horváth et al., 2019).

Parental effects, or transgenerational effects, unfold when the environmental experience by the parents affects the phenotype of their offspring, independent of genetic inheritance, and can be mediated by one or both parents, and are accordingly dubbed either maternal, paternal or parental effects (Marshall & Uller, 2007; Reddon, 2012). In many animals, the environmental variations experienced by parents are known to have critical effects on their progeny’ morphology, physiology, and mean behavioral traits (Tang et al., 2012; Ensminger et al., 2018; McDonald & Schwanz, 2018). Parents experiencing specific conditions can furnish information about these conditions to their offspring and by that way influence their phenotypic trajectories and behavior in adulthood (Bernardo, 1996). Adjusting their offspring personality, or increasing the offspring survival, is perceived as a beneficial strategy to augment parental fitness (Reddon, 2012). The influence of parents, particularly maternal impacts, on the personality of their offspring was documented across various animal taxa. However, most research has concentrated on stress-induced maternal effects related to food availability or composition, with a lack of studies addressing maternal effects stemming from various favorable food options that do not lead to critical deficiencies or nutritional imbalances. For example, work in rock lizards, Iberolacerta cyreni, shows that the boldness of the juveniles was affected by increasing vitamin D3 level in the maternal diet and corticosterone treatment early in life (Horváth et al., 2019). Harten et al. (2021) found that in the Egyptian fruit bat Rousettus aegyptiacus, a high cortisol level in the milk of urban-origin bat mothers influenced the personality in boldness and exploration of their new-born offspring. They were bolder, faster learning, but less exploratory than the rural-origin sibling. Another maternal effect was recorded by Petelle et al. (2017) that the offspring personality in boldness was affected by maternal age together with glucocorticoid metabolite levels (stress levels) in yellow-bellied marmots Marmota flaviventris. Offspring from older mothers with high level of glucocorticoid were consistently shier than offspring from younger mothers.

Here, we aimed to find out whether and how different favorable maternal diets can affect the offspring’ personality. We tested our hypothesis in plant-inhabiting, omnivorous predatory mites Amblyseius swirskii Athias-Henriot (Acari, Phytoseiidae). These mites are well suited models for our study questions because they can utilize diverse diets, including plant-associated herbivorous and predatory mites, small insects, and plant-derived substances such as pollen (Vangansbeke et al., 2016). Previous studies showed that A. swirskii are conducive to parental diet effects and have diet-mediated personalities. Parental, and even grandparental, diets may greatly influence the mean foraging behavior and cognition in A. swirskii offspring (Reichert et al., 2017; Schausberger & Rendon, 2022). Personal early life diet experiences can mediate the expression of adult personalities of A. swirskii (Nguyen & Schausberger, 2024). In this study, the mothers of the experimental predatory mites experienced, during egg production, one of three common diets, two-spotted spider mites Tetranychus urticae Koch (Acari, Tetranychidae), narrow-leaf cattail pollen Typha angustifolia L. (Poales, Typhaceae), or western flower thrips Frankliniella occidentalis Pergande (Thysanoptera, Thripidae). We then evaluated the influence of the maternal diet on the adult daughters and sons’ mean trait expression and repeatability in activity, exploration, and boldness.

## Materials and methods

### Study animals and rearing

Amblyseius swirskii used in experiments were randomly collected from a population reared in the laboratory, established by specimens collected on citrus trees in Cyprus, on acrylic platforms. Each platform consisted of an acrylic tile placed on a water-saturated foam block, delimited by tissue paper wrapped around the edges of the tile, inside a plastic box (20 x 20 x 5 cm) half-filled with tap water. Cotton tufts under coverslips (10 x 10 mm) served as shelters for the mites. To avoid escaping and contamination by other organisms, the plastic box containing the rearing platform was placed inside a plastic tray (40 x 30 x 8 cm), containing a shallow layer of soapy water, and covered by an acrylic case with a mesh-covered opening (10 cm Ø) on top for ventilation. Typha angustifolia pollen (Nutrimite, Biobest, BE) and mixed stages of T. urticae were added onto the platforms twice a week.

The laboratory population of T. urticae (green form) used as prey was reared on whole common bean plants, Phaseolus vulgaris var. Maxi, grown in a peat moss/sand mixture in 12 cm Ø pots. The laboratory population of thrips F. occidentalis used as prey was reared on green bean pods of P. vulgaris and T. angustifolia pollen inside a translucent plastic jar with a fine mesh cover for ventilation, stored in an air-conditioned room (23 ± 1 °C; 60 ± 5 % RH). Fresh bean pods and pollen were added to the jars once a week. To obtain first larval instars of thrips, adult females were randomly withdrawn from the rearing and transferred to a detached bean leaf, embedded upside down in a 1.5 % water agar solution in a closed Petri dish, for oviposition. After 24 h, the females were removed and the Petri dishes with thrips eggs inside the leaves were stored in a growth chamber at 23 ± 1 °C, 60 ± 5% relative humidity RH, and 16:8 h L:D photoperiod. First larval instars, hatching after 4 to 5 d, were used as prey during maternal conditioning.

### Maternal treatments and experimental animals

To assess maternal diet effects on offspring mean behavior and personality, we examined the behavior of adult A. swirskii females and males emerging from mothers that had been fed on either cattail pollen, spider mites, or thrips during egg production. To generate the mothers of the experimental animals, gravid females of A. swirskii were randomly collected from the rearing and placed, in batches of 15 females, for oviposition on new acrylic platforms (constructed as described above) harboring pollen. Predatory mite eggs were collected from the platforms and distributed, in batches of 10 eggs, to small acrylic arenas, each equipped with a cotton tuft shelter and cattail pollen. The predatory mites were left on the arenas until the females were adult and mated. Mated females were distributed to three types of new small acrylic arenas (5 to 7 females/arena), each equipped with a cotton tuft shelter and harboring either cattail pollen, mixed stages of two-spotted spider mites, or live and dead 1^st^ instar larvae of thrips; some thrips larvae were killed by deep-freezing before transfer to ease feeding by the predatory mite females (Nguyen & Schausberger, 2024). Eggs produced by the predatory females during the first two days of feeding on their respective diet were discarded to avoid any carryover effect from the previous diet on egg production and provisioning (Schausberger & Rendon, 2022). From the 3rd day onwards, predatory mite eggs were collected daily from each of the three diet-specific arenas and rinsed with de-ionized water for 30 s to remove any diet cues on the chorion. Eggs from the same maternal diet treatment were transferred to a new small acrylic arena harboring T. angustifolia pollen and left there until reaching adulthood and mating. Mated males (sons) and females (daughters) were used as experimental animals. Before starting the behavioral assays, each experimental individual was singly transferred to a small leaf disc (10 mm Ø), resting on top of a water agar column inside a closed breeding dish (50 x 15 mm), with a mesh-covered ventilation opening in the lid, half-filled with water, and provided with T. angustifolia pollen. These leaf discs were considered the home discs of each experimental individual, where they were kept before and in between the behavioral assays until the end of the experiment. Pollen on the discs was provided ad libitum and replenished as needed. All arenas and leaf discs were stored in a climate chamber at 23 ± 1 °C, 60 ± 5 % RH, and 16:8 h L:D.

### Behavioral assays

The mean trait expressions and personalities of adult A. swirskii females and males were assessed in three of the five canonical personality categories proposed by Réale et al. (2007). Personalities in exploration and boldness were evaluated by subjecting each individual to three successive tests for each trait. Personalities in activity were assessed when residing on their familiar home discs in between the assays for exploration and boldness. Throughout the behavioral assays, we also quantified reproduction of the experimental females by recording the number of eggs produced in the test arenas and on the home discs. The sample sizes were 98 individuals for boldness (34, 31, 33 for pollen, spider mites, thrips), 92 for activity (32, 30, 30 for pollen, spider mites, thrips), and 89 for exploration (31, 29, 29 for pollen, spider mites, thrips).

Personality in activity was assessed by observing each predator on their home disc three times within one hour (0, 30 and 60 min), each on five days in between the exploration and boldness assays. At each spot observation the individual was scored as either moving (ambulating) or being stationary.

Personality in boldness was evaluated by the response of the experimental animals to intraguild predator cues in three sequential tests. Each boldness test was staged using closed T-shaped acrylic mazes. Each T-maze was composed of two large cavities and one small cavity that were connected with each other by a T-shaped aisle. The bottom of the maze was closed by a fine mesh and on the upper side by a removable glass slide fixed by two foldback clips (Schausberger & Hoffmann 2008). The two large cavities, located at either end of the horizontal aisle of the T, were sites with (risky) and without (safe) intraguild predator cues; the small cavity was located at the bottom end of the vertical aisle of the T-maze and used as the release site of the experimental animal. We used cues of three different intraguild predators of A. swirskii, namely Orius laevigatus, A. andersoni, and Neoseiulus californicus in the three boldness tests. To generate intraguild predator cues inside one of the two large cavities, one adult intraguild predator female was placed inside one of the large cavities for 12 h before starting the boldness test (the aisle leading to the other cavities was blocked), to leave her traces, such as metabolic waste products and/or footprints, inside the cavity (considered as a risky site). The other large cavity without the predator traces was considered as a safe site. After removing the intraguild predator and the aisle blockage, the boldness test started by releasing an experimental female or male A.swirskii in the small cavity. In each boldness test, the T-shaped mazes were monitored immediately after release of A. swirskii and then every 20 min over 2 h to record their residence (distance to risky site) and activity (stationary or moving).

Personality in exploration was assessed by exposing the predators to a battery of three tests, offering a novel diet type in the first test, novel objects in the second test, and a novel leaf disc in the third test. In the first test, an arrangement of two plastic discs (15 mm Ø) connected by a non-fragrant wax bridge (28 mm long) on a wet filter paper resting on a water-saturated foam block (6 x 6 x 4 cm) inside a small plastic box (10 x 10 x 6 cm). One disc harbored T. angustifolia pollen, which was familiar to the predators, while the other disc harbored tomato pollen, Solanum lycopersicum, which was novel to the predators. The experimental animals were singly transferred to the familiar disc of the test arrangement and stayed there for 5 min for acclimatization. During these 5 min, a moist tissue strip was placed on the wax bridge to prevent crossing the bridge during acclimatization. To start the 1^st^ test, the moist tissue strip was removed and the activity and position of the experimental animal was videotaped for 5 min, using a digital stereo-microscope (Leica DMS1000). Videos were then watched to determine (i) the dispersal latency, i.e., the time elapsed until the predator first moved onto the wax bridge, and (ii) the object contact latency, i.e., the time elapsed until the predator reached the 2nd disc harboring the novel tomato pollen. In the 2nd test, an open-field arena (Antunes & Biala, 2012) was used to evaluate the propensity of the predators to explore novel objects. Each open-field arena consisted of a light-grey or milky white acrylic tile (60 x 60 x 3 mm) resting on a water-saturated foam block (6 x 6 x 4 cm) inside a plastic box (10 x 10 x 6 cm) half-filled with tap water. Moist tissue paper was wrapped around the edges of the tile to delimit the experimental arena. The four corners of the open-field arena were equipped with piles of paper pieces with different shapes (rectangular, circular, triangular) and colors (pink, light green, white), and a pile of cut plastic hairs (6 to 7 mm long). A small plastic washer (5 and 2.5 mm outer and inner Ø, 1 mm high), with some T. angustifolia pollen grains inside, was placed in the center of the arena to serve as shelter and release point for the predatory mites. Before starting the test, a single adult predatory mite was placed inside the plastic washer and the washer was closed by a coverslip. After a 5-min acclimatization period, the coverslip was removed, and the movement of the predator was videotaped for 5 min using a digital stereo microscope (Leica DMS1000). From the videos, the exploration propensity was quantified by determining (i) the dispersal latency, i.e. the time elapsed until the predator left the washer and moved away from it by at least one body length, and (ii) the object contact latency, i.e. the time elapsed until the predator reached, and touched, one of the novel objects in the corners of the arena. The 3rd test was conducted using a closed acrylic cage consisting of two circular cavities (15 mm Ø) connected by an aisle (20 mm long), all drilled into an acrylic plate (3 mm thick) closed at the bottom by a fine mesh and on the upper side by a removable glass slide fixed by foldback clips (Schausberger & Hoffmann, 2008). A small leaf disc of the familiar common bean (10 mm Ø) was placed inside one cavity to represent a familiar environment for the predator. The other cavity was loaded with a novel leaf disc (10 mm Ø) of blushing philodendron (Philodendron erubescens). To allow the predators to acclimatize to the familiar cavity, the aisle was blocked with a paper plug for 5 min. The test was started by removing the paper plug and subsequently the activity and position of the predator was recorded for 5 min. Exploration behavior was quantified by (i) the dispersal latency, i.e., the time elapsed until the predator first moved into the aisle leading to the novel leaf disc, and (ii) the object contact latency, i.e., time elapsed until the predator reached the second cavity with the novel leaf disc. If the experimental animal did not move away from the release site or did not reach the novel object during the experiment, a ceiling time of twice the observation time (10 min) was used in analysis of the dispersal and contact latencies.

### Statistical analysis

All statistical analyses were performed using R (R Core Team, 2024) and RStudio (Posit Team, 2024). All figures were created using the ggplot2 package (Wickham, 2016). If needed, dependent variables were log-transformed to normalize the data before analysis.

Separate generalized linear models (GLMs) and linear mixed-effects models (LMERs) were used to test the effect of maternal diet (pollen, spider mites or thrips) on the offspring’ mean activity on their home disc (proportion moving in five tests; LMER, normal distribution), mean boldness (distance to risky site; LMER, normal distribution), mean exploration (dispersal and object contact latencies; LMERs, normal distribution), and mean activity during the boldness tests (proportion moving in three tests; GLM, normal distribution). These models included the maternal diet, the offspring sex and their interaction as fixed factors. Depending on model requirements, either test or predator identity were added as random factors (lmer4 package; Bates et al., 2015). The effect of maternal diet on the mean number of eggs laid by daughters was analyzed by GLM (Poisson distribution). Each analysis started with the full model and then removed all non-significant interaction terms in a stepwise fashion until arriving at the most parsimonious model.

Personalities in activity, boldness, and exploration were assessed by intraclass correlation coefficients (ICCs; two-way random, consistency, average measure) across all three maternal diet treatments and two sexes, and separately for each diet treatment, for each sex and the combination of these factors, using the irr package (Gamer et al., 2019). Among-and within-individual variances were scrutinized to pinpoint the causes of differences in ICCs.

To evaluate the within-group composition of personality types as influenced by the maternal diet and offspring sex, each experimental animal was assigned a personality score. Personality scores in activity ranged from 0 to 4, with scores based on consistency in the proportion of time moving over the five tests (supplementary table 1). Personality scores in boldness ranged from 0 to 6, based on the consistency in the distance kept to the risky site over the three tests (supplementary table 1). Personality in exploration ranged from 0 to 9 using the consistency in dispersal latency or object contact latency over the three tests (supplementary table 1). Separate GLMs (Poisson distribution) were used to analyze the influence of maternal diet and offspring sex on the personality composition in activity, boldness, and exploration.

The link between the fitness of the females (reproduction) and their personality scores was assessed by linear and quadratic regressions of the number of eggs laid by each female on the personality scores in activity, boldness, and exploration.

## Results

### Population means

The maternal diet had a significant effect on mean activity of the predators both in their home cages (GLM; 1Z² = 8.12, P = 0.02) and during the boldness tests (1Z² = 6.61, P = 0.04) as well as on mean boldness (LMER; 1Z² = 5.91, P = 0.05), but not on mean exploration propensity (LMER, both parameters; 1Z² < 0.70, P > 0.71) (Fig. 1). Offspring from spider mite-fed mothers were the most active when they were in their home cage, while offspring from pollen-fed mothers were the least active during the boldness tests. Daughters were more active than sons both in their home cage (LMER; 1Z² = 19.42, P = 1.05e-05) and during the boldness tests (1Z² = 6.64, P = 0.01). Daughters and sons did not differ in mean boldness, as measured in residence relative to the risky site (LMER: 1Z² = 0.08, P = 0.77) and mean exploration (LMER: 1Z² < 1.28, P > 0.26). Maternal diet did not influence egg production of daughters (GLM; 1Z² = 3.82, P = 0.15).

**Figure 1.**
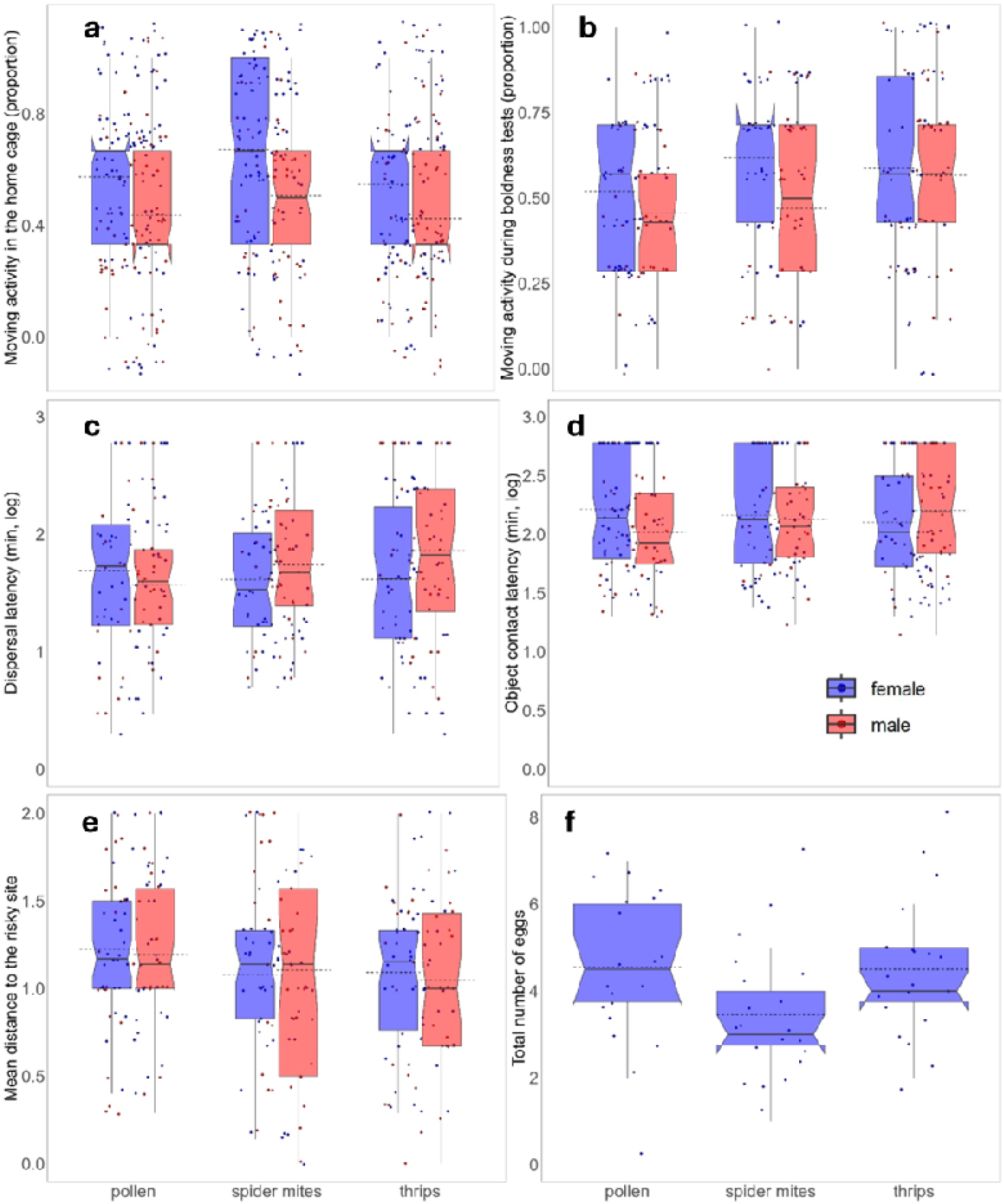
Mean activity in the home cage (a) and during the boldness tests (b), exploration (dispersal latency - c, object contact latency – d), and boldness (e) of Amblyseius swirskii males and females, and total number of eggs (f) produced by A. swirskii females in dependence of the maternal diet. Horizontal solid lines are the medians, dashed lines are the means, boxes are the 1^st^ and 3^rd^ quartiles, whiskers represent the standard errors (±1), dots indicate individual data points.

### Personality expression (ICCs)

Activity was moderately repeatable pooled across maternal treatments, both when the predators were in their home cages and during the boldness tests (Table 1). While both daughters and sons were repeatable in activity in their home cage, only sons were repeatable in activity during the boldness tests (Table 1). Regarding the maternal treatment-specific ICCs in activity, only offspring, especially daughters, from thrips-fed mothers were repeatable in activity in their home cage (Table 1). This was mediated by decreased within-individual variance and increased among-individual variance relative to the other treatments. Offspring from spider mite-fed mothers showed moderate repeatability in activity when they were in their home cage, which was mainly the case in sons (due to increased among-individual variance). In contrast, only daughters were repeatable in activity during the boldness tests (due to decreased within-individual variance) (Table 1). Offspring from pollen-fed mothers were marginally significantly repeatable in activity in their home cage whereas sons showed moderate repeatability in activity during the boldness tests (due to decreased within-individual variance) (Table 1).

**Table 1.** Behavioral repeatability (personality), quantified by intraclass correlation coefficients (ICC; [2,2]), of adult Amblyseius swirskii females and males as affected by the maternal diets. Significant ICCs and P-values (P ≤ 0.05) are highlighted in bold.

| Personality dimension | Treatment | *Pooled* |  | *Female* |  | *Male* |  |
| --- | --- | --- | --- | --- | --- | --- | --- |
|  |  | ICC | P-value | ICC | P-value | ICC | P-value |
| Activity | Pooled | **0.46** | **3.42e-05** | **0.30** | **0.04** | **0.48** | **0.004** |
|  | Pollen | 0.33 | 0.06 | 0.25 | 0.19 | 0.05 | 0.42 |
|  | Spider mites | **0.38** | **0.04** | -0.19 | 0.64 | **0.69** | **0.004** |
|  | Thrips | **0.59** | **0.0004** | **0.50** | **0.02** | **0.63** | **0.01** |
| Boldness - activity | Pooled | **0.30** | **0.02** | 0.21 | 0.16 | **0.39** | **0.04** |
|  | Pollen | 0.24 | 0.18 | -0.09 | 0.57 | **0.58** | **0.01** |
|  | Spider mites | **0.53** | **0.007** | **0.59** | **0.01** | 0.30 | 0.24 |
|  | Thrips | -0.03 | 0.52 | -0.06 | 0.54 | 0.17 | 0.34 |
| Boldness - residence | Pooled | 0.11 | 0.26 | 0.18 | 0.19 | 0.07 | 0.40 |
|  | Pollen | 0.37 | 0.06 | 0.31 | 0.18 | 0.46 | 0.10 |
|  | Spider mites | -0.27 | 0.75 | 0.25 | 0.23 | -1.34 | 0.90 |
|  | Thrips | 8.14e-05 | 0.49 | -0.42 | 0.77 | 0.25 | 0.27 |
| Exploration – dispersal latency | Pooled | 0.07 | 0.34 | 0.009 | 0.47 | 0.15 | 0.27 |
|  | Pollen | 0.19 | 0.24 | 0.35 | 0.13 | -0.30 | 0.68 |
|  | Spider mites | 0.31 | 0.12 | 0.30 | 0.17 | 0.21 | 0.31 |
|  | Thrips | -0.20 | 0.71 | -0.70 | 0.89 | 0.15 | 0.35 |
| Exploration – object contact latency | Pooled | -0.23 | 0.87 | -0.45 | 0.94 | 0.05 | 0.42 |
|  | Pollen | -0.19 | 0.70 | -0.89 | 0.93 | 0.004 | 0.48 |
|  | Spider mites | 0.005 | 0.48 | 0.02 | 0.46 | -0.01 | 0.48 |
|  | Thrips | -0.41 | 0.86 | -0.67 | 0.89 | -0.11 | 0.56 |

Boldness and exploration pooled across maternal treatments were not repeatable. The treatment-specific ICCs showed that only offspring from pollen-fed mothers were marginally significantly repeatable in boldness (Table 1). None of the offspring from the three maternal diet treatments showed repeatability in either dispersal latency or object contact latency (Table 1).

### Within-group personality composition and fitness

Maternal diet had a marginally significant effect on the offspring’ personality composition in boldness (GLM; 1² = 5.25, P = 0.07), with maternal thrips diet shifting the boldness scores up (Fig. 2b). Maternal diet had no effect on the offspring’ personality composition in activity (GLM; 1² = 2.02, P = 0.36) or exploration (GLM; 1² = 4.29, P = 0.12) (Fig. 2acd). Personality composition in object contact latency varied between sons and daughters (GLM; 1² = 12.37, P < 0.001; Fig. 2d), which was not the case for dispersal latency (GLM; 1² = 1.77, P = 0.18; Fig. 2c). Sons comprised more little exploratory types than daughters in object contact latency (Fig. 2d). However, this was only true in offspring from thrips-and spider mite-fed mothers whereas the opposite was the case in offspring from pollen-fed mothers (GLM; maternal diet x offspring sex: 1² = 10.78, P = 0.005). Personality composition in boldness was similar between sons and daughters (GLM; 1² = 0.0006, P = 0.98), whereas there was a marginal difference between the sexes in the personality composition in activity (GLM; 1² = 3.05, P = 0.08). Regarding object contact latency, sons from thrips-fed mothers were the least exploratory type compared to female offspring.

**Figure 2.**
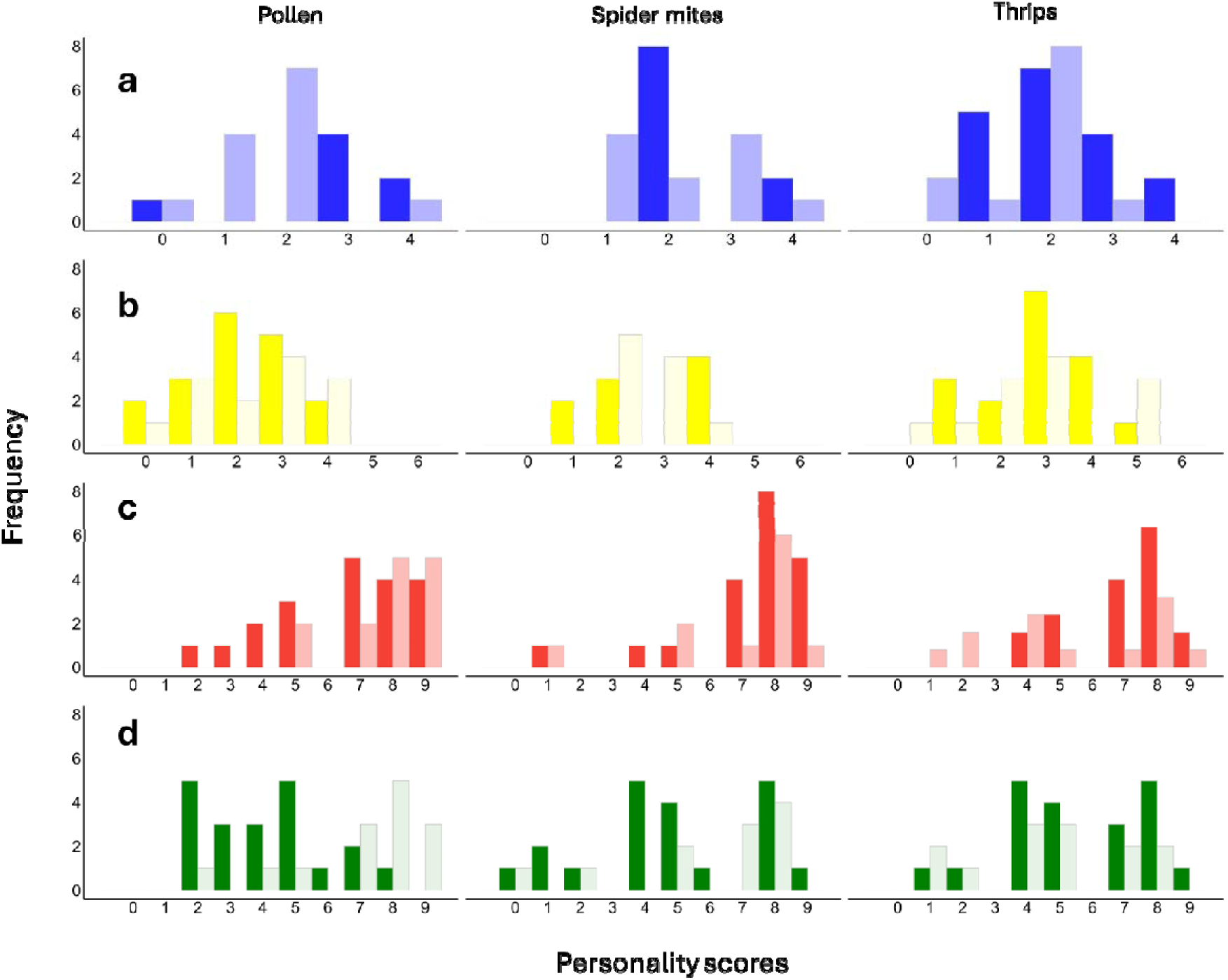
Histograms of personality types in activity (a), boldness (b), and exploration (dispersal latency - c; object contact latency - d), of Amblyseius swirskii females (dark bars) and males (light bars) in dependence of the maternal diet. Marginal personality scores indicate highly consistent types, whereas intermediate values indicate more plastic types (see supplementary table 1 for characterization of personality types).

Across maternal diet treatments, only the personality types in activity correlated with egg production, with consistently very active and consistently inactive personality types producing more eggs than inconsistent, flexible types (quadratic regression: r² = 0.14, P = 0.02; Fig. 3a). Regarding exploration, maternal spider mite diet produced offspring’ personality types in object contact latency, which correlated marginally positively with egg production (quadratic regression: r² = 0.26, P = 0.07): inconsistent flexible types in object contact latency produced the fewest eggs (Fig. 3d).

**Figure 3.**
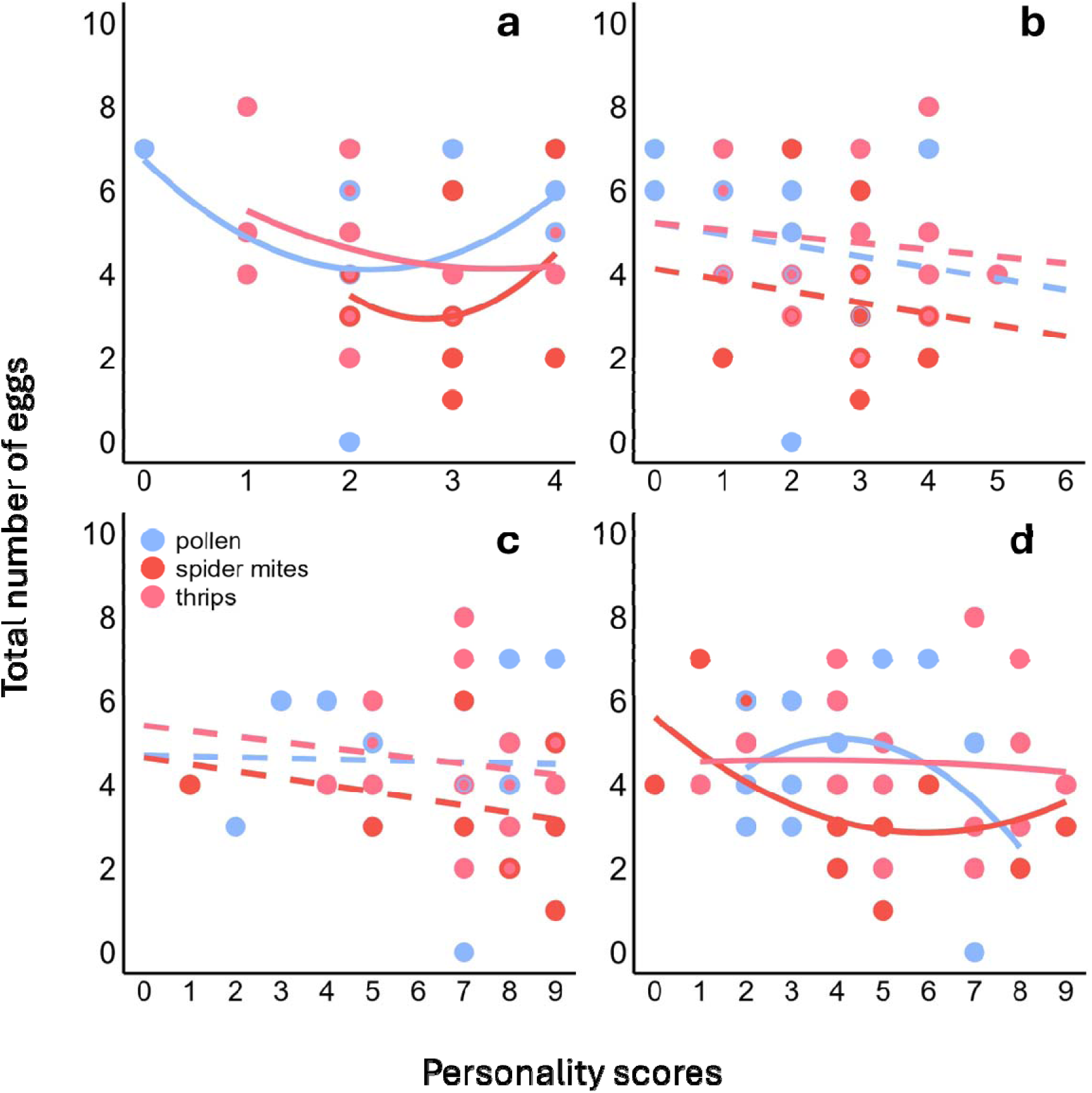
Total number of eggs regressed (linear, quadratic) on the personality scores in activity (a), and boldness (b), exploration (dispersal latency - c; object contact latency - d) of Amblyseius swirskii females in dependence of the maternal diet. Marginal personality scores indicate highly consistent types, whereas intermediate values indicate more plastic types (see supplementary table 1 for characterization of personality types). Solid and broken lines indicate significant and non-significant regressions.

## Discussion

Our study documents trans-generational impacts of maternal diet on mean behavioral traits as well as personality expression of adult offspring in plant-inhabiting predatory mites A. swirskii. Our experimental design allows to unambiguously assign the observed effects to mothers since the experimental diets were only administered after development and mating (all fathers in all maternal treatments were fed the same pollen diet). Maternal diet influenced offspring’ mean activity and boldness, which effects differed between sons and daughters in activity but not boldness. Offspring activity was the only personality trait that was repeatable across maternal treatments. In two of three personality traits, repeatability varied with maternal diet and offspring sex. The composition of personality types in boldness changed slightly with maternal diet, with maternal thrips diet shifting the offspring’ boldness scores up. Among sons, there were more consistently highly exploratory and consistently little active personality types than among daughters. Consistently highly and little active daughters produced more eggs across maternal diet treatments. Consistently little or highly exploratory daughters from spider mite-fed mothers produced more eggs than flexible daughters.

### Population means

The offspring from prey-fed mothers were more active than those from pollen-fed mothers. It is perceivable that spider mites and thrips would exert a similar influence as animal diets provide elevated levels of suitable nutrients to primarily carnivorous predators, compared to vegetarian diets such as pollen (Lundgren, 2009; Delisle et al., 2015). Accordingly, spider mite-and thrips-fed mothers likely provided their offspring with higher initial energy reserves than pollen-fed mothers. Availability and composition of nutrients in the yolk during embryonic development can be important proximate factors mediating maternal effects (Deeming & Ferguson, 1991; Stewart & Ecay, 2010). However, from hatching onwards, all offspring received completely the same pollen diet, providing ample time and opportunity to equalize any presumable initial differences; still, metabolic memory could have carried initial differences over to adulthood (Somer & Thummel, 2014). In any case, prey-fed mothers produced offspring with higher mean activity in familiar environments. Also, during the boldness assays, offspring from pollen-fed mothers were the least active, while offspring from thrips-and spider mite-fed mothers maintained similarly high activity. These results are analogous to those by Seiter & Schausberger (2016), who found that A.swirskii emerging from spider-mite fed parents were more active than those from pollen-fed parents. The pollen diet presumably indicates a steady, low-stress environment to mothers, which then program their offspring to maintain reduced activity as a little energy-consuming strategy. Nonetheless, these differences did not translate into differences in mean exploration propensities, which underscores that movement activity in familiar environments is distinct from activity during the boldness and exploration tests.

### Personality expression (ICCs)

Of the three personality traits evaluated, only activity was consistently repeatable across treatments, yet repeatability varied strongly among maternal diet treatments and between sons and daughters. Boldness was only repeatable in particular maternal diet treatments. Parental effects are anticipated to generate behavioral diversity and individual niche specialization within a population, facilitating flexible and rapid adjustment to changing environments, such as varying occurrence and availability of different diet options (Mousseau & Fox, 1998; Proulx & Teotónio, 2017). No previous studies exist that compared the effects of various nutritionally favorable maternal diets on the development of offspring personalities. Previous research primarily examined the impact of variations in availability of single diet components on mean behavioral trait expression and animal personality. Horváth et al. (2019) observed that maternal diet supplementation with vitamin D₃ affected the activity-related personality traits of offspring in rock lizards, Iberolacerta cyreni. In our experiments, thrips prey exhibited the most pronounced effects among the three maternal diet treatments. Thrips are difficult to capture (e.g. Schausberger et al., 2018), and maternal experience of thrips appears to promote the development of two distinct personality types in activity among offspring. This was similar when A. swirskii experienced thrips in early life (Nguyen & Schausberger, 2024). Also, the fact the exploration was only little repeatable across maternal diets aligns with previous studies on the effects of early-life diet experiences in A. swirskii (Nguyen & Schausberger, 2024). Nonetheless, personal early life experience of thrips prey generated highly repeatable personalities in exploration (Nguyen & Schausberger, 2024), which differed from the effects of maternal thrips diet on offspring observed in this study. This suggests that maternal and early-life experiences do not always and not necessarily act in the same direction, as shown in another predatory mite, Phytoseiulus persimilis (Nguyen & Schausberger, 2025), and that thrips prey has a stronger and persistent impact on predator behavior if it is personally experienced. Offspring from pollen-fed mothers were the only group among the three maternal diet treatments that were (marginally significantly) repeatable in boldness. Conversely, in rock lizards, vitamin D_3_ of the maternal diets did not affect offspring’ boldness repeatability (Horváth et al., 2019). On average, however, offspring from mothers supplied with vitamin D3 were bolder than those from control mothers. Harten et al. (2021) observed that urban fruit bats with high cortisol level in their milk, relative to rural fruit bat mothers, heightened the risk-taking tendency and decreased the exploration propensity of rural foster offspring, as compared to those raised by their biological mothers.

### Within-group personality composition and fitness

Our results indicate maternally-mediated changes in the distribution and composition of personality types in boldness. Specifically, maternal thrips diet shifted the offspring’ personality types in boldness up, which is similar to the effects of early-life intraguild predation risk observed by Schausberger et al. (2024). This suggests that mothers inform their offspring about foraging-associated risks, and that the offspring then adjust their personalities accordingly. Thrips is only a risky prey for juvenile but not adult predatory mites, which is similar to the risk posed by intraguild predators used in the boldness tests. It thus pays for adult offspring to not shy away from sites that no longer pose a risk to them, no matter whether in foraging or predation. In foraging contexts, maternal information transfer allows offspring to preferentially go for the type of maternally-experienced prey (Bilkó et al., 1994; Peralta Quesada & Schausberger, 2012). While there was no main effect of maternal diet on within-group personality composition in activity and exploration, sons and daughters responded differently to maternal diets. Among offspring from spider mite-and thrips-fed mothers, there were more cautious exploratory personality types in sons than daughters. This may be linked to the much smaller body size of males, which rendes them more vulnerable to risk and weaker in overwhelming prey. The opposite was the case in sons and daughters from pollen-fed mothers. Exploration typically reflects a trade-off between the benefits of acquiring new information and the potential costs of higher risk (Wolf et al., 2007) and this trade-off is inevitably linked to sex-specific acrivities and body size.

Personality traits such as activity, aggressiveness, exploration, sociability and boldness are assumed to have implications to fitness (e.g., Wolf et al. 2007; Biro & Stamps, 2008). For example, more exploratory personalities are more likely to discover new resources, which often enhances developmental speed and increases reproductive output (McCowan et al., 2014; Roth et al., 2021; Nguyen & Schausberger, 2025). Our experiments revealed convex correlations between egg production and personality scores in activity (across maternal treatments) and exploration (in offspring from spider mite-fed mothers). These results indicate that the maternal diet can mediate the adaptive significance of offspring’ personality expression. To our knowledge, no previous study has examined the correlation between within-group personality composition and reproduction as affected by parental effects.

## Conclusions

There exist only very few studies addressing maternal and/or paternal effects on animal personality, especially related to food intake. Apart from showing that maternal diet affects mean behavioral trait expression of offspring (Horváth et al., 2019; Stamps & Bell, 2021; Schausberger & Rendon, 2022; Nguyen & Schausberger, 2025), our study demonstrates that maternal diet during egg production can also have long-lasting impacts on offspring’ behavioral repeatability. This finding highlights the importance to consider transgenerational effects and parental diet history in studies on animal personality (Reddon, 2012). Proximate factors include epigenetic changes, embryonic learning, and differences in maternal and/or offspring’ (through maternal provisioning) initial metabolic or nutritional state having long-lasting (condition transfer or carry-over) effects into offspring’ adulthood (Reddon, 2012; Somer & Thummel, 2014; Peralta Quesada & Schausberger, 2012; Sih et al., 2015; Bonduriansky & Crean, 2018; Schausberger & Rendon, 2022). However, regarding nutritional state, there was ample time throughout ontogeny (same food for all offspring from the egg onwards) to balance any initial maternally-mediated differences. Differences could have also been mediated or strengthened by interactions during development in groups with similarly-programmed individuals (individuals of a given maternal treatment grew up together), including mating among those individuals. Recent studies suggest that the mating partners can have decisive effects on adult females’ personalities (Monestier & Bell, 2023; Schausberger et al., 2024; Schausberger & Nguyen, 2025). In our study, the mothers received the experimental diets only during internal production of the eggs giving rise to the experimental animals. Before that, all mothers and fathers had been fed on the same pollen diet. This procedure excludes paternal diet influences and suggests that the mothers’ response to short-term diet changes allows to flexibly diversify their offspring’ personalities and individual niches throughout the oviposition period. Diversification in mean behavior and personality, alongside lacking differences in mean egg production among daughters from the three maternal diet treatments, should mitigate inter-individual conflicts and facilitate the local and regional co-existence of inter-and intra-generationally adjusted personalities and foraging phenotypes (Nguyen & Schausberger, 2024, 2025; this study).

## Author contributions

THN and PS conceived the study and designed the experiments; THN conducted the experiments, analyzed the data, and wrote the first draft of the manuscript; PS provided resources, supervised the work and contributed to writing; THN and PS acquired funding.

## Acknowledgments

This work was financially supported by the Austrian Science Fund (FWF; P 33787-B to PS). THN was supported by a Dimitrov fellowship from the Austrian Academy of Sciences (ÖAW). Mustafa Altintas is thanked for helping with rearing the mites and plants.

