## Supplementary table 1 for "Maternal diet exerts sex-specific effects on offspring’ personalities in predatory mites"

**Supplementary table.** Categorization of personalities in boldness, activity, and exploration.

| **Personality score in boldness** | **Mean distance from the risky site in 3 tests** |
| --- | --- |
| 6 | ≤0.67, ≤0.67, ≤0.67 |
| 5 | ≤0.67, ≤0.67, 1.34>x>0.67 |
| 4 | ≤0.67, ≤0.67, ≥1.34/≤0.67, 1.34>x>0.67, 1.34>x>0.67 |
| 3 | ≤0.67, 1.34>x>0.67, ≥1.34/1.34>x>0.67, 1.34>x>0.67,1.34>x>0.67 |
| 2 | 1.34>x>0.67, 1.34>x>0.67, >1.34/≤0.67, ≥1.34, ≥1.34 |
| 1 | ≥1.34, ≥1.34, 1.34>x>0.67 |
| 0 | ≥1.34, ≥1.34, ≥1.34 |
| **Activity** | **Percentage of time moving in 5 tests** |
| 0 | <0.67, <0.67, <0.67, <0.67, <0.67 |
| 1 | <0.67, <0.67, <0.67, <0.67, >0.33 |
| 2 | <0.33, <0.33, >0.67, >0.67, >0.67/<0.33, <0.33, <0.33, >0.67, >0.67 |
| 3 | <0.33, <0.33, <0.33, <0.33, >0.67 |
| 4 | <0.33, <0.33, <0.33, <0.33, <0.33 |
| **Exploration** | **Dispersal latency or object contact latency** |
| 0 | 600,600, 600^1^ |
| 1 | 600, 600, > x̅ |
| 2 | 600, 600, < x̅ |
| 3 | 600, > x̅, > x̅ |
| 4 | 600, < x̅, > x̅ |
| 5 | 600, < x̅, < x̅ |
| 6 | > x̅, > x̅, > x̅ |
| 7 | > x̅, > x̅, < x̅ |
| 8 | >x̅, < x̅, < x̅ |
| 9 | < x̅, < x̅, < x̅ |

^1^ for the future time estimated in case individual did not move out or checked new object
